# Sinking particle export within and beneath the euphotic zone in the eastern Indian Ocean

**DOI:** 10.1101/2025.08.14.670365

**Authors:** Michael R. Stukel, Tristan Biard, Moira Décima, Christian K. Fender, Opeyemi Kehinde, Thomas B. Kelly, Sven A. Kranz, Manon Laget, Michael R. Landry, Natalia Yingling

## Abstract

The eastern Indian Ocean is substantially under sampled with respect to the biological carbon pump – the suite of processes that transport the carbon fixed by phytoplankton into the deeper ocean. Using sediment traps and other ecosystem measurements, we quantified sinking organic matter flux and investigated the characteristics of sinking particles in waters overlying the Argo Abyssal Plain directly downstream of the Indonesian Throughflow off northwest Australia. Carbon export from the euphotic zone averaged 7.0 mmol C m^-2^ d^-1^, which equated to an average export efficiency (export / net primary production) of 0.17. Sinking particle flux within the euphotic zone (beneath the mixed layer, but above the deep chlorophyll maximum) averaged slightly higher than flux at the base of the euphotic zone, suggesting that the deep euphotic zone was a depth stratum of net particle remineralization. Carbon flux attenuation continued into the twilight zone with a transfer efficiency (export at euphotic depth + 100m / export at euphotic depth) of 0.62 and an average Martin’s *b*-value of 1.1. Within the euphotic zone, fresh phytoplankton (chlorophyll associated with sinking particles, possibly contained within appendicularian houses) were an important component of sinking particles, but beneath the euphotic zone the fecal pellets of herbivorous zooplankton (phaeopigments) were more important. Changes in carbon and nitrogen isotopic composition with depth further reflected remineralization processes occurring as particles sank. We show similarities with biological carbon pump functioning in a similar semi-enclosed oligotrophic marginal sea, the Gulf of Mexico, including net remineralization across the deep chlorophyll maximum.

**Submitted to: Deep-sea Research II**

**Highlights:** Despite low productivity, export efficiency was 17% of primary production

Flux attenuation beneath the euphotic zone (EZ) was low for a tropical region

Sinking particle flux from the upper to lower EZ exceeded export from lower EZ

The deep EZ was a stratum of net particle remineralization (and net heterotrophy)

## 1. Introduction

Net primary production (NPP) by phytoplankton decreases the partial pressure of CO_2_ in the upper ocean, leading to CO_2_ flux from the atmosphere into the oceans. However, most of the fixed carbon is rapidly consumed and recycled in the surface ocean with only a portion transported into the deeper ocean by a suite of processes referred to as the biological carbon pump (BCP) (Martin *et al*., 1987; Buesseler and Boyd, 2009; Boyd *et al*., 2019). Globally, the BCP is estimated to account for 5 – 13 Pg C yr^-1^ of flux into the deep ocean (Henson *et al*., 2011; Laws *et al*., 2011; Siegel *et al*., 2014; Henson *et al*., 2022), although uncertainty is substantial and the BCP is highly variable at multiple spatial and temporal scales (Puigcorbé *et al*., 2017; Stukel *et al*., 2017; Smith *et al*., 2018).

The BCP includes multiple transport mechanisms, including active vertical migrations of zooplankton and nekton and passive transport of organic matter within subducted water (Carlson *et al*., 1994; Steinberg *et al*., 2000; Davison *et al*., 2013; Omand *et al*., 2015). However, the BCP is typically dominated by the carbon export of sinking particles (Nowicki *et al*., 2022; Stukel *et al*., 2022a; Stukel *et al*., 2023) involving processes that package smaller cells into aggregates and fecal pellets that can sink at speeds of hundreds of meters per day (Passow *et al*., 2001; McDonnell and Buesseler, 2010; Turner, 2015). The variety of ecological and biogeochemical processes involved, and their variability in time and space, complicate attempts to predict future changes in the BCP and marine carbon sequestration (Laufkötter *et al*., 2016; Henson et al., 2022).

Relative to the other major oceans, the Indian Ocean (IO) is substantially under sampled with respect to BCP-relevant processes (Hood *et al*., 2024a). It is a heterogeneous ocean with regions that are both net sources and net sinks for carbon dioxide (Bates *et al*., 2006; Ghosh *et al*., 2024; Hood *et al*., 2024b). The Arabian Sea, perhaps the most well-studied IO region, is strongly monsoon-influenced with high productivity during the southwest monsoon but with temporal variability at other scales and substantial spatial variability in productivity-limiting processes (Kumar *et al*., 2000; Barber *et al*., 2001; Marra and Barber, 2005). Substantial variability in sinking carbon export flux of this region follows from its diversity of productivity and trophic conditions (Buesseler *et al*., 1998; Lee *et al*., 1998; Honjo *et al*., 1999; Sarma *et al*., 2003; Rixen *et al*., 2019). A north-south transect from the Arabian Sea (17°N to 12°S) showed low export efficiency of 1.0 – 4.4% of primary production leaving the euphotic zone during boreal spring, its most oligotrophic state (Subha Anand *et al*., 2018). In contrast, few carbon export measurements are available for the eastern IO, although the low export efficiency during oligotrophic conditions in the Arabian Sea is likely a reasonable expectation for similarly oligotrophic conditions in other IO regions.

This study focuses on export processes overlying the Argo Abyssal Plain, a deep oceanic basin (hereafter, Argo Basin) in the eastern Indian Ocean (Fig. 1) that was investigated as part of the Second International Indian Ocean Expedition (IIOE-2). The Argo Basin is bounded on the south by northwestern Australia and on the north by the Indonesian archipelago. Remote sensing products show that it is a tropical, oligotrophic system with low surface chlorophyll concentration and suggest low net primary production throughout the year (Kehinde *et al*., 2023; Hood et al., 2024b). The Argo Basin is the only known spawning site of Southern Bluefin tuna (Davis *et al*., 1990) and biogeochemically interesting as the terminus of the Indonesian Throughflow, the only low-latitude connection between the Pacific and Indian Oceans (Domingues *et al*., 2007; Sprintall *et al*., 2024). On the BLOOFINZ-IO (Bluefin Larvae in Oligotrophic Ocean Foodwebs, Investigations of Nutrients to Zooplankton – Indian Ocean, Landry *et al*., this issue-a) Cruise in Jan – Mar 2021, we measured the magnitude and characteristics of sinking particles within and beneath the euphotic zone of the Argo Basin. We use these results to quantify and test the expected inefficiency of the BCP in this oligotrophic spawning region and to investigate its ecological controls. We also compare dynamics of the deep chlorophyll maximum of the Argo region to another oligotrophic bluefin tuna spawning basin in the deepwater Gulf of Mexico that has many similar characteristics as a warm, deep-water, oligotrophic region and was sampled by similar methods as part of the BLOOFINZ-GoM Program (Gerard *et al*., 2022).

**Figure 1.**
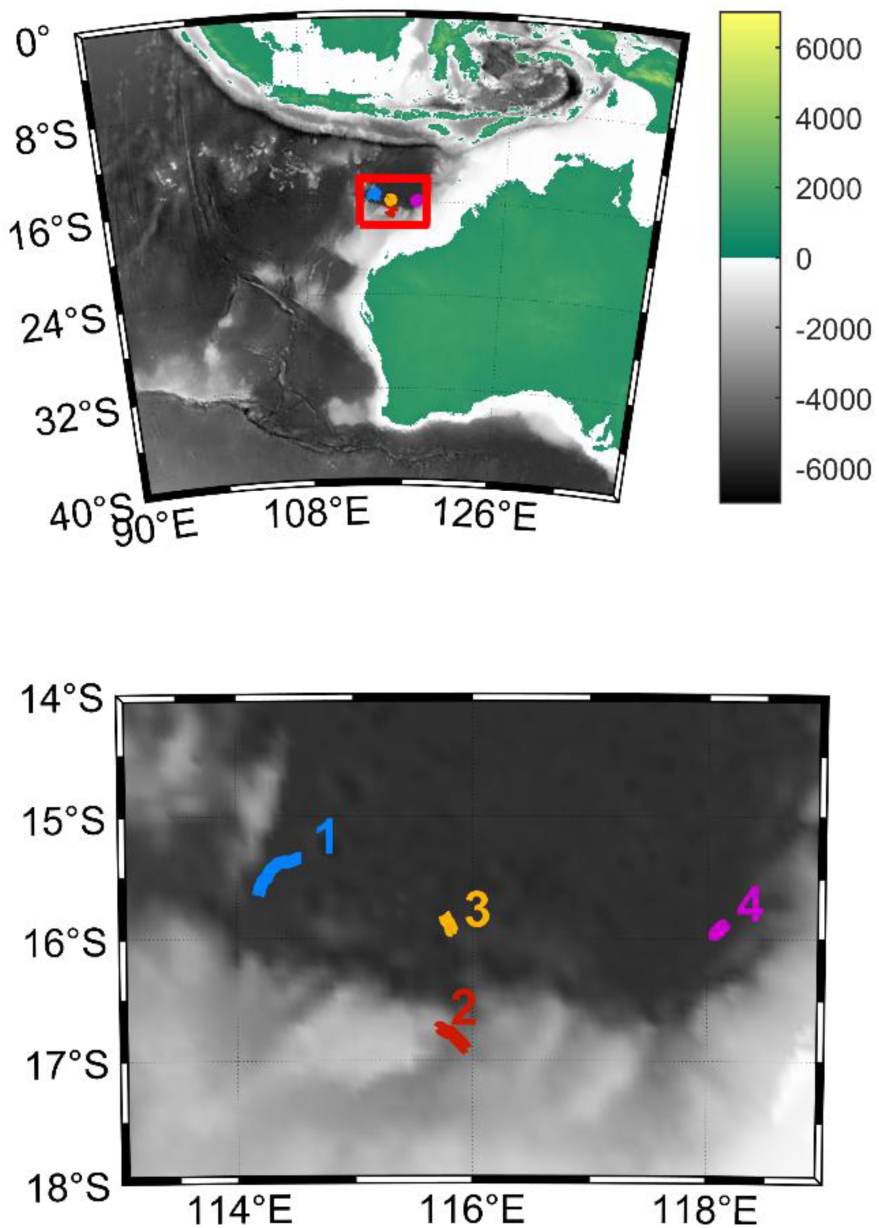
Map of the study region with the top panel showing the broader area and bottom panel the study region itself, depicted with a red box on the top panel. Color represents elevation in both. Lagrangian trajectories of sediment traps are shown and labelled by cycle.

## 2. Methods

### 2.1. Experimental design and water column measurements

Samples were collected in the Argo Basin in the eastern Indian Ocean as part of the BLOOFINZ-IO cruise from January – March 2022 (Landry et al., this issue-a). As with the companion study in the Gulf of Mexico (BLOOFINZ-GoM, Gerard et al., 2022), this cruise was organized around Lagrangian experiments (hereafter “cycles”). Water parcels of interest with respect to physical and biotic characteristics were identified through a combination of satellite imagery, ship underway flow-through data, and repeated net sampling (Landry et al., this issue-a). Water parcels were then followed using a pair of floating arrays. Both arrays included a satellite-enabled surface float tethered to a 3×1-m holey sock drogue (Landry *et al*., 2009). One array (described further below) also included sediment trap crosspieces for collecting sinking particles. The other array (“incubation array”) had stainless steel rings integrated into the line at fixed depths below the surface float for attaching mesh bags containing experimental bottles that were incubated in situ for 24-hr periods to measure net primary production and other ecosystem rates (Landry et al., 2009).

Lagrangian experiments lasted from 3.3 to 4.3 days. At 02:00 local time each day, a CTD with attached Niskin rosette was deployed to measure vertical profiles of temperature, salinity, fluorescence, and oxygen and to collect samples from 6 depths (from the near surface to the DCM depth) for nutrients, chlorophyll a (Chl*a*), particulate organic matter (POM), and NPP measurements. Chl*a* and POM were also measured at additional deeper depths. Nutrient samples were filtered through a 0.1-µm Acropak cartridge filter and stored frozen until analysis for nitrate+nitrite, nitrite, ammonium, phosphate, and silicic acid using an autoanalyzer at the Scripps Institution of Oceanography, Ocean Data Facility. Samples for Chl*a* and phaeopigments were filtered through glass fiber fine (GF/F) filters at low vacuum pressure and measured on a Turner 10AU fluorometer using the acidification method after 24-h acetone extraction at -20 °C. POM samples (4-L) were filtered through pre-combusted GF/F filters, which were dried until analysis on land. Filters were then acidified to remove inorganic carbon, packed into pre-combusted tin capsules and analyzed for carbon, nitrogen, and isotopes at the University of California, Davis, Stable Isotope Facility.

Phytoplankton community composition was quantified using a combination of flow cytometry for picoplankton (<2 µm) and epifluorescence microscopy for nano- and microplankton (Yingling *et al*., this issue). Samples for phytoplankton community composition were collected daily from the same depths as chlorophyll, nutrients, and net primary production. Picophytoplankton biomass was estimated from FCM abundances assuming a cellular carbon content of 36, 101 and 359 fg C cell^-1^ for *Prochlorococcus*, *Synechococcus*, and picoeukaryotes, respectively (Yingling et al., this issue). The biomass of nano- and microphytoplankton was estimated using biovolume measurements from epifluorescence microscopy and allometric carbon:volume relationships from Menden-Deuer & Lessard (2000).

Triplicate samples (plus a dark blank) for net primary production were gently transferred to polycarbonate bottles which were spiked with H^14^CO_3_^-^ and incubated at in situ depths on the incubation array for 24 h. After recovery, samples were filtered through GF/F filters and ^14^C radioactivity was determined with a liquid scintillation counter (Kranz *et al*., this issue). An Underwater Vision Profiler 6 (UVP6) was attached to the CTD-Niskin rosette and used to measure vertical profiles of the particle volume-size spectrum (Picheral *et al*., 2022). We restricted analysis of this data to particles (including aggregates) with sizes ranging from 51 – 1290 µm. Below 51 µm, pixel-level noise made the data untrustworthy; above 1290 µm the sampled volume was too low for reasonable particle count estimates.

### 2.2. Sediment traps

VERTEX-style sediment trap crosspieces (Knauer *et al*., 1979) were attached to the sediment trap array at four depths: beneath the mixed layer (52-62 m), near the base of the euphotic zone as estimated from fluorescence profiles (116 – 127 m), and in the mesopelagic zone at ∼220 and ∼420 m depth. Aqualogger pressure-temperature sensors were used to measure actual depths of trap deployments. Twelve particle interceptor tubes (7-cm inner diameter, 8:1 aspect ratio, topped with a baffle constructed of 14 smaller tubes that were tapered on the top) were attached to each crosspiece at the three shallowest depths (16 tubes at the deepest depth). Tubes were deployed with a seawater brine made from 0.1-µm seawater amended with 50 g L^-1^ NaCl and formaldehyde (0.4%, final concentration) and left for 3.3 to 4.3 days. After recovery, overlying water was immediately removed from each tube by gentle suction. Samples were then filtered through a 100-µm Nitex filter and the filter was inspected at 25X magnification under a stereomicroscope to remove zooplankton “swimmers”. Contents of filters (i.e., non-swimmer sample) were then washed back into the filtrate (<100-µm portion of the sample). Three tubes per depth were filtered through pre-combusted GF/F filters for carbon, nitrogen, and isotope samples (analyzed as described above), and 50-mL subsamples were taken from three separate tubes and analyzed for Chl*a* and phaeopigments as described above.

### 2.3 BLOOFINZ-GoM comparison cruise

Nearly identical experimental methods were used as part of the BLOOFINZ-GoM companion cruises in an oligotrophic area of the Gulf of Mexico (Gerard et al., 2022; Stukel *et al*., 2022b). This region is similar to the Argo Basin as a semi-enclosed basin with a deepwater, oligotrophic interior that is a primary spawning site for bluefin tuna larvae (Southern Bluefin tuna in Argo Basin, Atlantic Bluefin tuna in Gulf of Mexico). BLOOFINZ-GoM cruises were conducted in May 2017 (BLOOFINZ-GoM Cycles 1 – 3) and May 2018 (BLOOFINZ-GoM Cycles 4 – 5). Slight methodological differences between cruises include the absence of a deep (>400 m) sediment trap cross-piece (Stukel *et al*., 2021), the use of H^13^CO_3_ ^-^ instead of H^14^CO_3_ ^-^ to measure NPP (Yingling *et al*., 2022), and lack of a UVP6 in the GoM.

## 3. Results

### 3.1. Oceanographic and water column conditions in the Argo Basin

The surface ocean was warm and strongly stratified throughout the cruise. The deepest cycle-average mixed layer depth (defined as a 0.01 kg m^-3^ increase in potential density relative to the average potential density of the upper 5 m) was 30.5 m for Cycle 1, which was conducted shortly after a storm moved through the region. Even for this cycle, however, surface temperatures exceeded 28°C (Fig. 2a). For the other cycles, cycle-average mixed layer depth was always shallower than 10 m (Fig. 2b), while sea surface temperatures (calculated at 5-m depth) were 28.8, 29.8, and 30.5 °C for Cycles 2 – 4, respectively.

**Figure 2.**
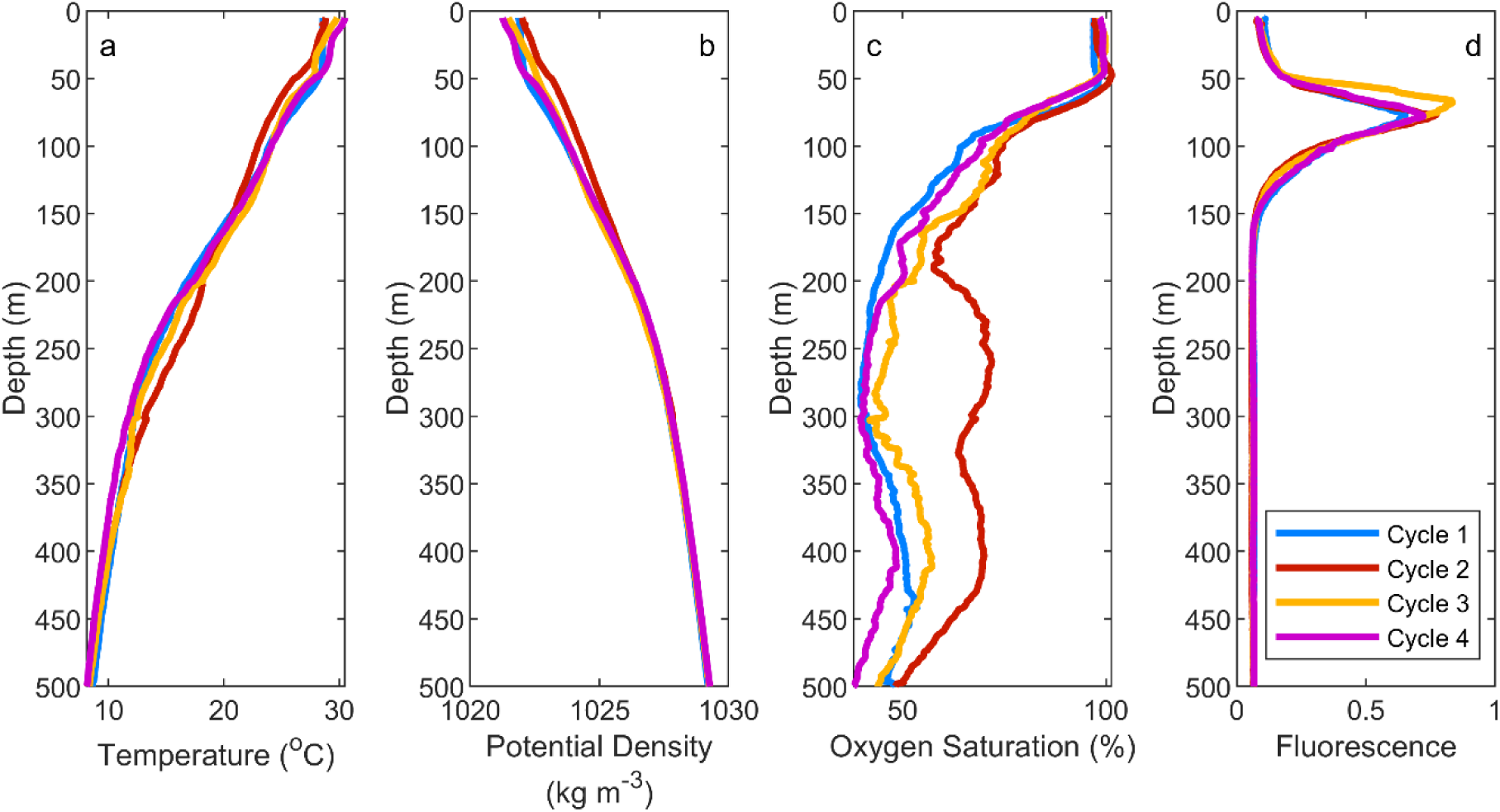
Cycle average vertical profiles of (a) temperature, (b) potential density, (c) oxygen saturation, and (d) fluorescence (proxy for chlorophyll).

Warm surface temperatures and stratified conditions were paired with strong deep chlorophyll maxima (DCM) that ranged from 67 (Cycle 3) to 75 m (Cycle 1) depth (Figs. 2d, 3a). Chl*a* concentrations were much higher at the DCM (cycle averages ranged from 0.29 to 0.56 mg Chl*a* m^-3^) than at the surface (0.07 to 0.09 mg Chl *a* m^-3^). Nitrate concentration was consistently low above 50 m with an average of 0.016 µmol L^-1^ across all samples and a maximum measured value (above 50 m) of 0.12 µmol L^-1^. Nitracline depths (defined as the depth at which nitrate first exceeded 1 µmol L^-1^) were near the DCM, with cycle-average nitracline depths ranging from 66 to 78 m. The cycle-average depths of the euphotic zone (here, defined as the depth of the 0.1% light level) ranged from 121 to 127 m. Oxygen concentrations were near saturation with the atmosphere in the upper ∼50 m, but decreased rapidly with depth beneath 50 m and typically were ∼50% saturation at ∼200-m depth (Fig. 2c). In contrast to chlorophyll, particulate organic carbon (POC) concentrations varied only weakly with depth in the upper 250 m of the water column. POC increased slightly with depth from the surface to the DCM and then decreased mostly monotonically with depth in deeper waters. Despite these trends, POC typically varied within a small range of 2 – 5 mmol C m^-3^ throughout the epipelagic (Fig. 3b). At most depths, Cycle 4 had lower Chl*a* and POC concentrations than the other cycles, although differences were relatively small. These slight differences may reflect temporal variation across the system as nutrients injected by the storm in the beginning of the cruise were consumed over the following weeks.

**Figure 3.**
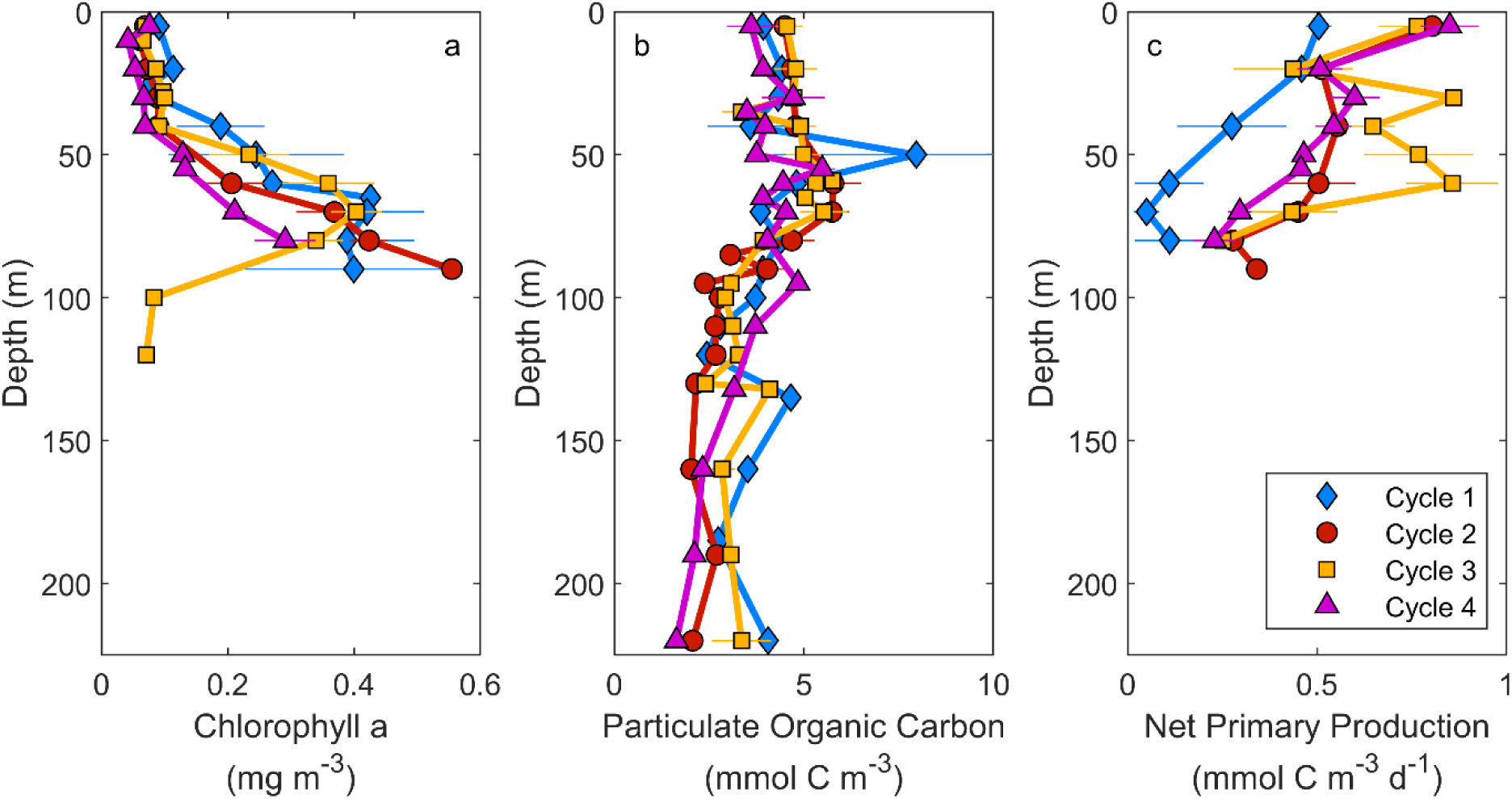
Vertical profiles of (a) Chlorophyll *a*, (b) Particulate Organic Carbon, (c) Net Primary Production. Average values and standard error for each Lagrangian cycle are shown.

Using a UVP6, we obtained estimates of particle abundances and volumes (including aggregates and living organisms) as a function of particle diameter and depth (Fig. 4). Large particles and aggregates (100 – 1000 µm diameter) were typically most abundant in the DCM depth range. Large particles were also abundant in the shallowest depth stratum (0 – 5 m), although it is likely that images in this depth range were affected by bubbles forming near the ocean surface. Subsurface large particle maxima were most pronounced during Cycles 2 and 3. During Cycle 1, total particle volume only increased ∼2-fold from the surface to the depth of the DCM, and large particle abundance integrated over the euphotic zone was substantially lower than on the other cycles. During Cycle 4, large particles had similar abundance at the deep chlorophyll maximum to Cycles 2 and 3, but large particle abundance was substantially greater in the depth range of ∼20 – 50 m during Cycle 4, relative to other cycles. This increased abundance of large particles during the later portions of the cruise may reflect an increased abundance of appendicularians (filter-feeding pelagic tunicates that produce large mucous feeding mesh “houses”), which were found to increase in abundance by more than a factor of 2 during the cruise (Swalethorp *et al*., this issue). Beneath the DCM, large particle abundance decreased rapidly with depth for all cycles.

**Figure 4.**
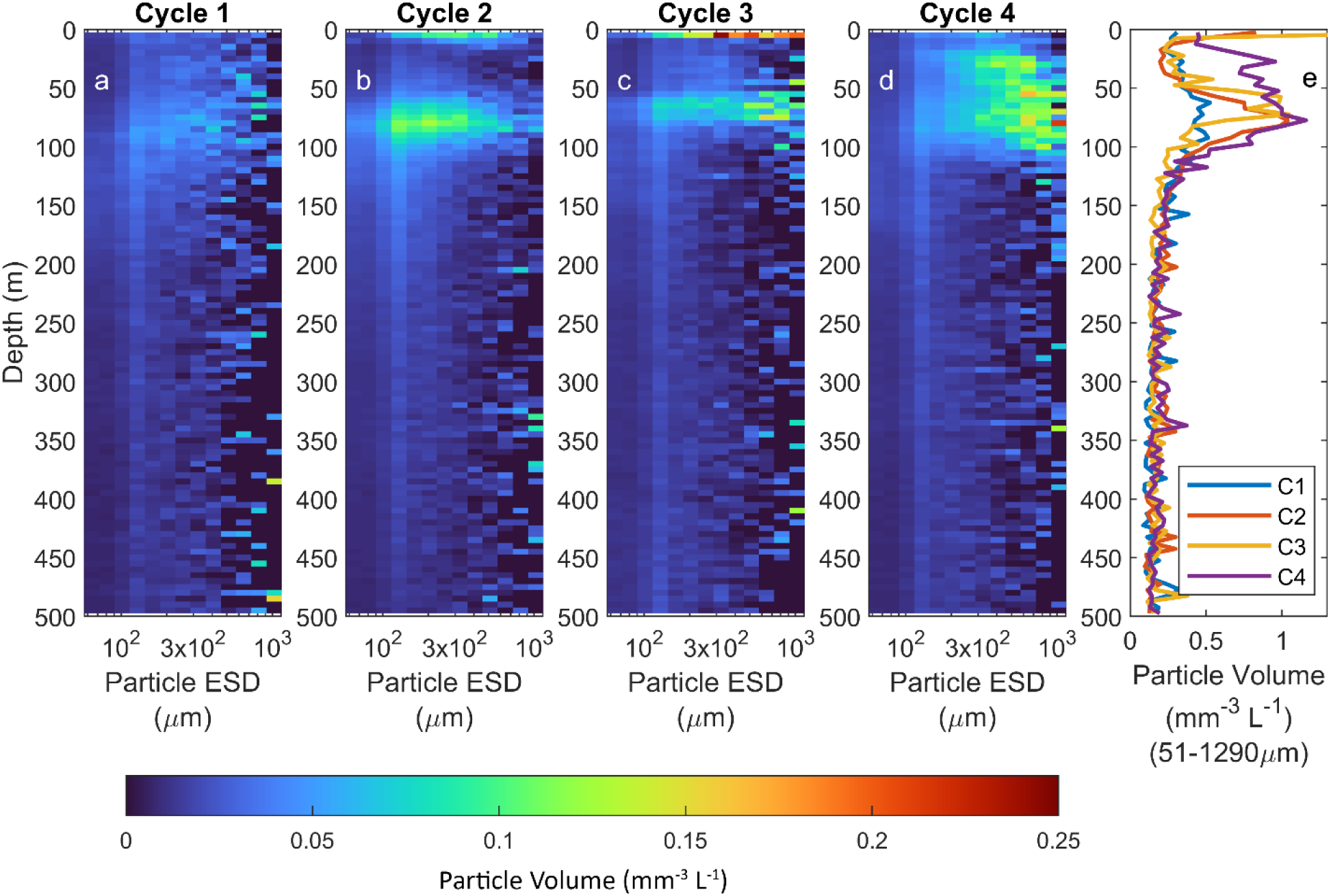
Large particle volume estimated with the Underwater Vision Profiler 6 (UVP6) as functions of equivalent spherical diameter (ESD, x-axis in panels a – d) and depth. a – d represent mean particle volumes (mm^-3^ L^-1^) from CTD casts for Cycles 1 – 4, respectively. Panel e shows summed particle volume across all size classes (from 51 to 1290 µm)

Vertically integrated NPP measured by H^14^CO_3_^-^ uptake was relatively low, as expected, for oligotrophic waters, with an average across all profiles of 37 ± 3 mmol C m^-2^ d^-1^. NPP was substantially lower on Cycle 1 (22.5 ± 5.5 mmol C m^-2^ d^-1^) than the other cycles, while Cycle 3 had slightly higher productivity (51.9 ± 4.1 mmol C m^-2^ d^-1^) than Cycles 2 and 4 (46.6 ± 3.8 and 41.3 ± 2.3 mmol C m^-2^ d^-1^, respectively). Low NPP on Cycle 1 may have been tied to light limitation; Cycle 1 had the lowest average surface photosynthetically-active radiation. NPP generally decreased monotonically with depth, although Cycle 3 exhibited little change with depth in the upper 60 m. Despite strong chlorophyll maxima at 67 to 75 m depth, NPP was typically much lower at the DCM than in the mixed layer.

### 3.2. Sinking particle flux in the Argo Basin

Organic carbon fluxes into sediment traps near the base of the euphotic zone (trap depth of 116 to 127 m) averaged 7.3 mmol C m^-2^ d^-1^ and ranged from 3.7 to 10.4 mmol C m^-2^ d^-1^ (Fig. 5a). Carbon fluxes during Cycles 1 and 2 were substantially lower than for Cycles 3 and 4. For traps placed at 52-62 m depth, below the mixed layer but above the DCM, carbon flux was actually higher on average (although not statistically different) than beneath the euphotic zone (mean = 8.1, range = 7.0 to 9.4 mmol C m^-2^ d^-1^). On Cycles 1 and 2, organic carbon flux decreased across the DCM, while on Cycles 3 and 4 organic carbon flux increased across the DCM. Beneath the euphotic zone, carbon flux decreased monotonically with depth, averaging 3.9 mmol C m^-2^ d^-1^ at 220 m (range 3.1 to 5.1 mmol C m^-2^ d^-1^) and 2.2 mmol C m^-2^ d^-1^ at 420 m (range 1.8 to 2.9 mmol C m^-2^ d^-1^). Combining data from all cycles, and using the 0.1% light level as the reference depth, a power law fit suggests an average Martin’s b-value of 1.1. This is fairly typical of global ocean b-value estimates (Marsay *et al*., 2015), suggesting moderate flux attenuation with depth, but notably lower than expected for a warm-water region (typical b of 1.2 - 1.6, Lamborg *et al*., 2008; Marsay et al., 2015). Sinking nitrogen fluxes mostly mirrored the patterns in sinking carbon (Fig. 5b). For Cycles 3 and 4, the C:N ratios of sinking particles remained relatively constant with depth (ranging from 7.2 to 10.3 mol:mol), while for Cycles 1 and 2 the C:N ratio increased with depth to an average of 15.5 (mol:mol) at 420 m.

**Figure 5.**
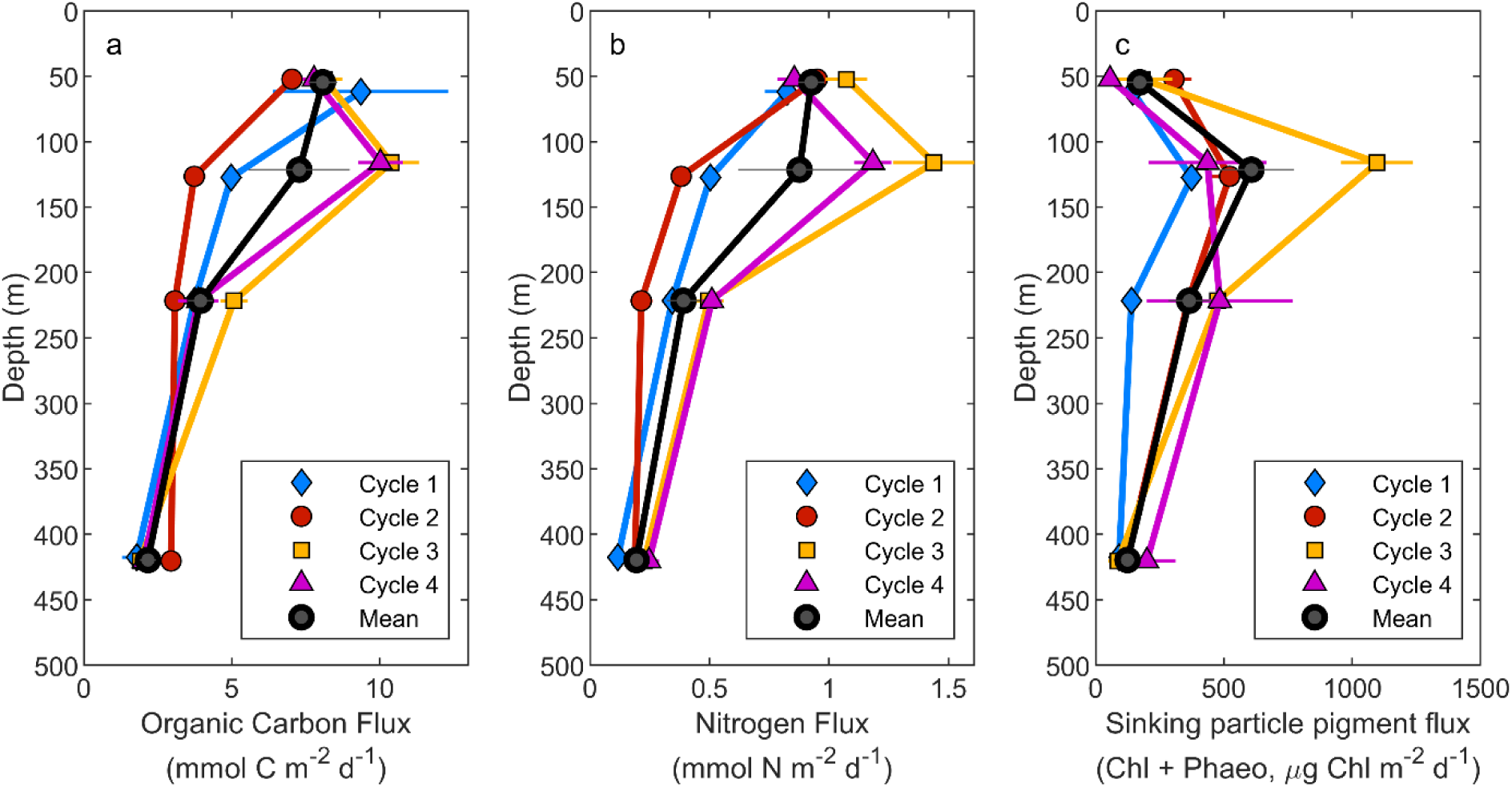
Sinking particle fluxes measured by sediment traps. (a) Organic Carbon Flux. (b) Nitrogen Flux. (c) Pigment Plux (sum of chlorophyll a and phaeopigments). Error bars are ± 1 standard error of the mean.

Carbon isotope values of sinking particles were lighter with depth, reflecting remineralization processes (Fig. 6b). This trend was consistent for all cycles and across all depths, except for an isotopically lighter sample at 200 m than at 420 m for Cycle 3. In comparison, nitrogen isotopes did not exhibit clear trends with depth. Nitrogen isotopes became isotopically heavier with depth for three cycles but showed no change or a slight decrease for Cycle 4, which was also isotopically lighter than the other cycles at shallow depths (Fig. 6c). Notably, suspended POM for Cycle 4 did not have similarly low δ^15^N as the sinking particles, with Cycle 2 having the lowest δ^15^N POM (Supp. Fig. 1). Looking across all samples, there was a strong inverse correlation between the δ^13^C and δ^15^N of sinking particles (Pearson’s ρ = -0.79, p = 2.8×10^-4^), but no similar correlation was found for suspended POM.

**Fig. 6.**
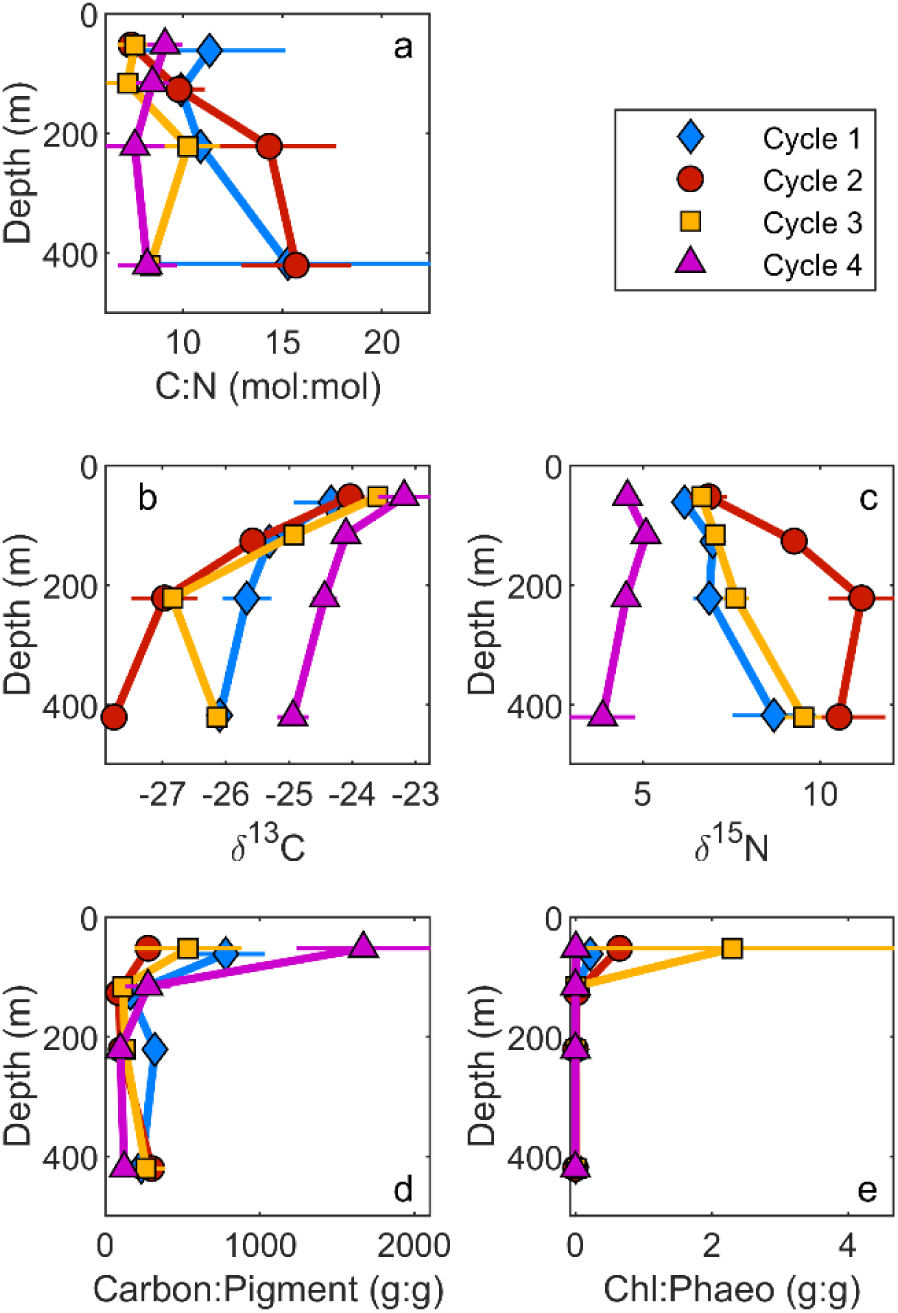
Characteristics of sinking particles. a) Organic Carbon:Nitrogen Ratio, b) ^13^Carbon Isotopic Values, c) ^15^Nitrogen Isotopic Values, d) Carbon:Pigment Ratio (pigment = chlorophyll *a* + phaeopigments), e) Chlorophyll *a*:Phaopigment Ratio. Means ± 1 standard errors are shown.

Total pigment (Chl*a* + phaeopigment) flux on sinking particles increased substantially from above the DCM (average of 172 µg pigment m^-2^ d^-1^) to beneath it (average of 607 µg pigment m^-2^ d^-1^, Fig. 4c). This increase in pigment flux was associated with a shift from a substantial flux of Chl*a* to predominantly phaeopigment flux. For Cycles 1 – 3, Chl*a* flux at 52–62 m ranged from 26 to 127 µg Chl*a* m^-2^ d^-1^. Beneath the euphotic zone (116–127 m), Chl*a* flux declined to 8.5 µg Chl*a* m^-2^ d^-1^ on Cycle 2 and was unmeasurable on all other cycles. In contrast, phaeopigment flux at 116–127 m increased to 374–1097 µg Chl*a* m^-2^ d^-1^. Because phaeopigments are generally indicative of fecal pellets produced by herbivorous zooplankton, this suggests a shift from sinking of uningested phytoplankton within the euphotic zone to primarily sinking fecal pellets exiting the euphotic zone. Phaeopigment flux then declined gradually with depth beneath the euphotic zone, with carbon:pigment ratios of sinking particles remaining relatively constant with depth (Fig. 6d). The decrease in carbon:total pigment flux within the euphotic zone likely reflects a change in the carbon:chl *a* ratio of phytoplankton with depth. Based on flow cytometry and epifluorescence microscopy-based estimates of phytoplankton carbon biomass (Yingling et al., this issue), phytoplankton carbon:chl *a* ratios averaged 130 g:g in the upper 50 m of the water column, and decreased to an average of 37 g:g in the 70–100 m depth range. If we assume that 130 g:g is representative of phytoplankton sinking into our shallowest sediment trap, then sinking phytoplankton flux ranged from 0.004 to 1.4 mmol C m^-2^ d^-1^ (with a mean of 0.74 mmol C m^-2^ d^-1^) and contributed up to 18% of sinking carbon above the DCM. These undigested sinking phytoplankton may have been attached to discarded appendicularian houses. These houses can be efficient mechanisms for concentrating small phytoplankton carbon into larger particles capable of relatively fast settling rates (Alldredge, 2005; Lombard and Kiorboe, 2010).

Beneath the euphotic zone, sinking carbon flux of uneaten phytoplankton was negligible. However, if we take 37 g:g as an estimate of the carbon:chl*a* of phytoplankton consumed by herbivorous zooplankton in the deep euphotic zone and that 70% of the ingested carbon is consumed while all of the chl*a* is converted to phaeopigments, then herbivore fecal pellets were responsible for 3.8 to 11.2 mmol C m^-2^ d^-1^ (with a mean of 6.2 mmol C m^-2^ d^-1^) and contributed to 45 to 141% of sinking carbon.

## 4. Discussion

### 4.1. The biological carbon pump in the Argo Basin and Gulf of Mexico

The oligotrophic central waters of the Gulf of Mexico is an interesting site for comparison with the Argo Basin, because it is also a semi-enclosed, tropical, deep-water basin and crucial spawning site for bluefin tuna (Atlantic Bluefin in the Gulf of Mexico, Southern Bluefin in the Argo Basin). Conditions encountered during the BLOOFINZ-GoM study were even more oligotrophic than those during the BLOOFINZ-IO cruise (Gerard et al., 2022). Nitracline and DCM depths during BLOOFINZ-GoM were deeper than in the Argo Basin (84 to 127 m and 78 to 137 m, respectively, in the GoM, Knapp *et al*., 2021; Landry *et al*., 2021, compared to 66 to 78 m and 67 to 75 m, respectively in the Argo Basin). Surface Chl*a* was similar to BLOOFINZ-IO (0.05 – 0.13 mg m^-3^) and *Prochlorococcus* dominated phytoplankton community biomass in both regions (Selph *et al*., 2021; Yingling et al., this issue). Sea surface temperatures were substantially cooler (∼26 °C) in the Gulf of Mexico than in the present study (28.8 – 30.5 °C).

Across the five BLOOFINZ-GoM cycles, NPP ranged from 24.3 – 29.3 mmol C m^-2^ d^-1^, which was lower than all but Cycle 1 of BLOOFINZ-IO (Fig. 7a). At other well-studied oligotrophic regions NPP typically averages ∼38 mmol C m^-2^ d^-1^; the range at the Hawaii Ocean Timeseries site in the North Pacific Subtropical Gyre is 26 - 49 mmol C m^-2^ d^-1^ and the range at the Bermuda Atlantic Time-Series site in the Sargasso Sea is 27 – 44 mmol C m^-2^ d^-1^ (Church *et al*., 2013). The Argo Basin thus exhibited fairly typical NPP during our cruise, while the Gulf of Mexico had slightly below average NPP, reflecting its extremely low levels of nutrients in the upper 100 m and very deep chlorophyll maxima. In contrast to those other oligotrophic sites, however, the Argo Basin and Gulf of Mexico had relatively high export efficiency. The EZ-ratio (ratio of export at the base of the euphotic zone to NPP) ranged from 0.085 to 0.23 for BLOOFINZ-IO and from 0.11 to 0.25 for BLOOFINZ-GoM (Fig 7b). These EZ-ratios, averaging 0.19 for BLOOFINZ-IO and 0.17 for BLOOFINZ-GoM, are substantially higher than typically expected for oligotrophic systems dominated by small phytoplankton (usually <10%, with a typical value of 5%, Buesseler and Boyd, 2009; Siegel et al., 2014; Karl *et al*., 2021; Stukel *et al*., 2024). Export is expected to be inefficient in these systems for multiple reasons: The supply of new nutrients necessary to support export (new production) is low (Eppley and Peterson, 1979; Harrison *et al*., 1987); the picoplankton that dominate the system are too small to sink individually (Smayda, 1970); and food-web pathways that convert phytoplankton into rapidly sinking zooplankton fecal pellets are typically long and inefficient (Michaels and Silver, 1988; Steinberg and Landry, 2017). These criteria for low export efficiency are all met in both regions: nutrient concentrations are too low to suggest substantial upwelled nitrate; *Prochlorococcus* dominates the phytoplankton biomass (Selph et al., 2021; Yingling et al., this issue); and food chains are long with substantial energy dissipation through the microbial loop (Landry et al., 2021; Stukel et al., 2022b; Stukel *et al*., this issue). High export efficiency is thus surprising, although it may be related to lateral transport from the adjacent productive coastal areas. Kelly et al. (2021) showed that all of the measured export flux for BLOOFINZ-GoM could be supported by lateral transport of POM (both living and dead) from the shelf region. Similarly, Kehinde et al. (2023) suggested that 32% (but with an uncertainty range of 10 - >100%) of export during BLOOFINZ-IO could be supported by lateral transport of POM. It is possible that this POM subsidy is additive to the normal low export efficiency of oligotrophic systems, although it is important to note that at the warm temperatures in the region, we should expect particles to be consumed, re-worked and/or recycled multiple times in the euphotic zone before sinking. Additionally, it is likely that the typical relationship between phytoplankton size and export efficiency was substantially modified by the observed importance of appendicularians (Swalethorp et al., this issue), which feed efficiently on picoplankton and produce large marine snow aggregates through their discarded mucus houses (Alldredge, 2005).

**Figure 7.**
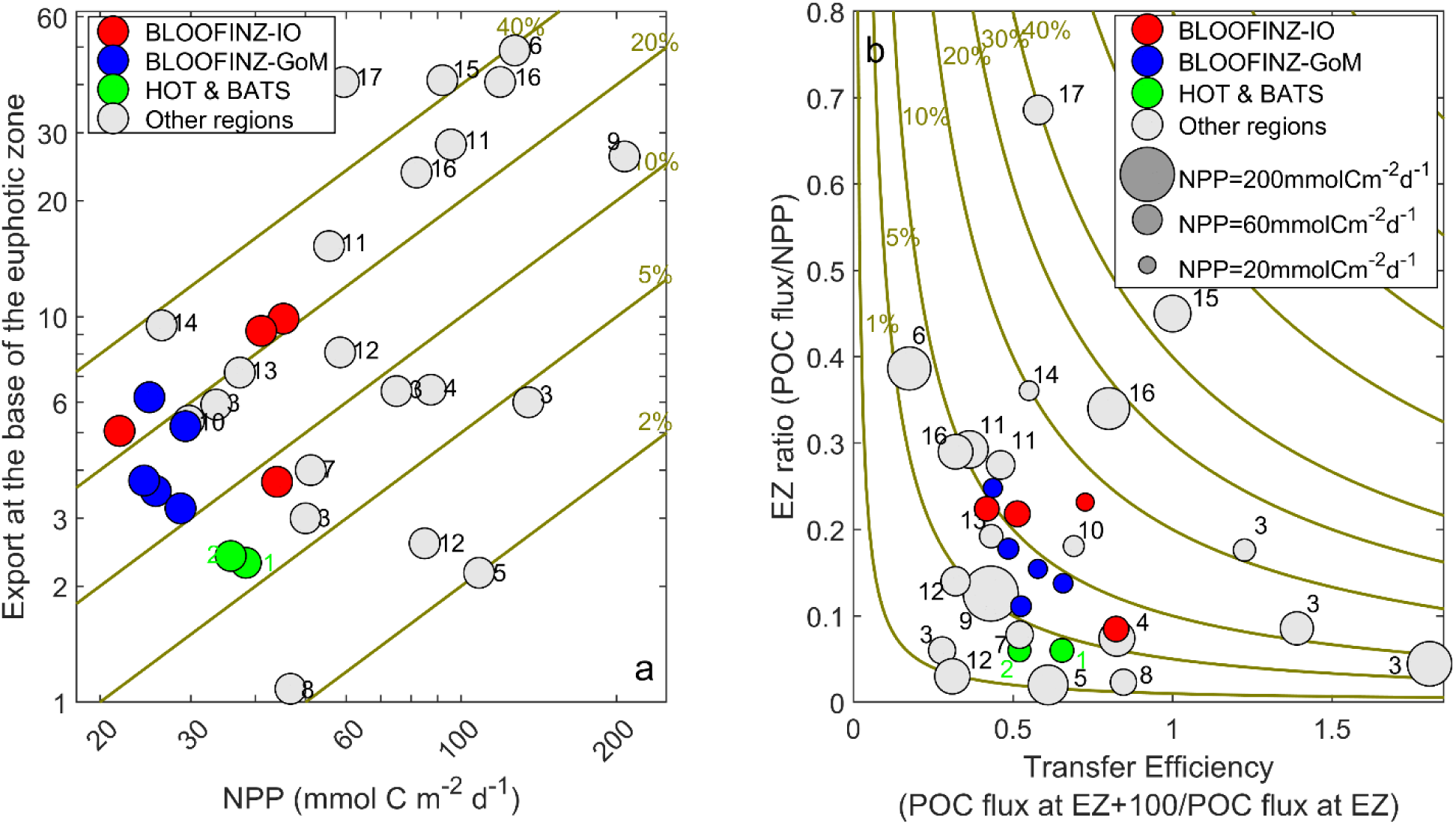
Comparisons of BLOOFINZ export to other regions. (a) shows sinking organic carbon flux at the base of the euphotic zone (0.1% light level) as a function of net primary production (NPP), with gold lines showing isopleths of constant EZ-ratio. (b) shows EZ-ratio plotted against transfer efficiency (T_100_ = export flux at euphotic depth + 100 m / export flux at euphotic depth). Gold lines show isopleths of constant percentage of NPP exported 100 m deeper than euphotic depth. In both panels red symbols are from Argo Basin, blue symbols are from Gulf of Mexico, and green symbols are from mean values from the warm-water, oligotrophic Hawaii Ocean Time-series (North Pacific Subtropical Gyre, ^1^(Church et al., 2013)) and Bermuda Atlantic Time-Series (Sargasso Sea, ^2^(Lomas *et al*., 2013)) Programs. With the exception of BLOOFINZ symbols, all results are means from multiple sediment trap deployments or ^234^Th profiles in a specific region. ^3^Sargasso Sea eddies (Buesseler *et al*., 2008a). ^4^Costa Rica Dome (Stukel *et al*., 2016). ^5^Equatorial Pacific (Bacon *et al*., 1996). ^6^Tropical South Pacific, shelf (Black *et al*., 2018). ^7^Tropical South Pacific, offshore (Black et al., 2018). ^8^Tropical South Pacific, gyre (Black et al., 2018). ^9^California Current Ecosystem, coastal upwelling (Stukel and Barbeau, 2020). ^10^California Current Ecosystem, offshore oligotrophic (Stukel and Barbeau, 2020). ^11^California Current Ecosystem, frontal regions (Krause *et al*., 2015; Stukel et al., 2017). ^12^Subarctic Pacific (Charette *et al*., 1999). ^13^Subarctic Pacific (Buesseler *et al*., 2008b). ^14^Spanish continental margin (Olli *et al*., 2001). ^15^North Atlantic Bloom (Buesseler *et al*., 1992). ^16^Southern Ocean (Buesseler *et al*., 2003). ^17^Barents Sea (Andreassen and Wassmann, 1998).

Particle transfer efficiency through the shallow twilight zone was moderate. The ratio of export at a depth horizon 100 m beneath the euphotic depth (0.1% light level) divided by export at the euphotic depth (T_100_ = flux transfer efficiency over 100 m) averaged 0.62 during BLOOFINZ-IO and 0.54 during BLOOFINZ-GoM (Fig. 7b). T_100_ was notably higher for Cycles 1 and 2 of BLOOFINZ-IO (0.73 and 0.82) than for Cycles 3 and 4 (0.51 and 0.42). These differences were driven by greater variability in export flux at the base of the euphotic zone than at other depths (Fig. 5a). The coefficient of variation (standard deviation / mean) for export flux interpolated to the base of the euphotic zone was 0.44, while the coefficient of variation was only 0.21 at a depth of ∼220 m and 0.25 at a depth of ∼420 m. Decreased variability in export flux in the mesopelagic relative to the base of the euphotic zone is likely a result of the “statistical funnel” effect (Siegel *et al*., 2008); the spatial imprint of surface production that contributes to export at a specific location grows with increasing depth as a result of the lag time introduced by slow particle settling speeds and rapid horizontal currents. Decreasing organic matter flux with depth was also associated with substantial shifts in the isotopic composition of sinking material. In both the GoM and Argo Basin, δ^15^N values generally increased with depth and δ^13^C values generally decreased with depth (Fig. 6 and Stukel et al., 2021). While the distinctly low δ^15^N for BLOOFINZ-IO Cycle 4 is likely indicative of recent diazotropy (Kranz et al., this issue), the strong correlation between δ^15^N and δ^13^C (Supp. Fig. 1f) suggests that remineralization processes may broadly drive patterns in both of these isotopes. Higher δ^15^N of sinking particles at deeper depths likely reflects isotopic fractionation as many metabolic processes preferentially utilize ^14^N (Post, 2002; Glibert *et al*., 2019). Conversely, lower δ^13^C suggests preferential utilization of organic compounds with comparatively high δ^13^C; more specifically, lipids have low δ^13^C and typically remain within sinking particles after amino acids have been remineralized (McConnaughey and McRoy, 1979; Wakeham and Lee, 2019).

During both BLOOFINZ-GoM and -IO studies, we deployed sediment traps in the euphotic zone at a depth of ∼50-60 m, beneath the mixed layer but above the DCM. Both studies showed that sinking carbon flux attenuation began within the euphotic zone. Organic carbon flux decreased from a mean of 8.1 mmol C m^-2^ d^-1^ at the shallowest depth to a mean of 7.0 mmol C m^-2^ d^-1^ at the base of the euphotic zone (0.1% light level) during BLOOFINZ-IO. It decreased even more strongly across this depth horizon during BLOOFINZ-GoM, declining from 6.4 to 4.4 mmol C m^-2^ d^-1^ (Supp. Fig. 2). This pattern of decreasing export flux through the deep euphotic zone was also more consistent in the Gulf of Mexico, declining during four out of five Lagrangian cycles. These changes in export flux with depth also included an increase in the pigment:carbon ratio and decrease in the Chl*a*:phaeopigment ratio for sinking particles in both BLOOFINZ-IO and BLOOFINZ-GoM, suggesting a switch from sinking phytoplankton within the euphotic zone to sinking fecal pellets at the base of the euphotic zone.

Particle size spectra can also be used to investigate processes relating to particle export and how it varies with depth. While early studies suggested that the particle (and aggregate) size spectra generated by the UVP could estimate sinking particle fluxes optically (Guidi *et al*., 2008), subsequent studies have shown that any such algorithms need to be regionally tuned (Iversen *et al*., 2010; Fender *et al*., 2019) and are likely not constant with depth (Fender et al., 2019). The variability in particle size-flux algorithms argues against treating UVP data as a reliable estimator of the absolute magnitude of sinking carbon flux. Nevertheless, comparison of sediment trap rates to UVP estimates of export flux from previously published algorithms allows us to investigate how the particle size – sinking speed relationship may change with depth (Supp. Fig. 3). Although none of the three algorithms provided accurate estimates of export, all showed similar patterns. Specifically, samples collected in the shallowest sediment traps had consistently lower particle fluxes than predicted by the algorithms, while the greatest underestimate by the UVP algorithms was for sediment traps located immediately beneath the euphotic zone (specifically on Cycles 3 and 4). This would suggest that particles and aggregates of a specific size were sinking more slowly within the euphotic zone (where high Chl*a*:C ratios indicated sinking phytoplankton that were loosely associated with aggregates or appendicularian houses) and more rapidly immediately beneath the euphotic zone (where high paheopigment:C ratios indicated sinking fecal pellets).

Zooplankton also contribute to carbon export by feeding in the surface ocean during the night and descending to mesopelagic depths during the day where their respiration leads to a net downward carbon transport (Longhurst *et al*., 1990; Steinberg et al., 2000). Decima et al. (this issue) measured day-night differences in zooplankton biomass and applied allometric scaling equations to estimate diel-vertical-migrating zooplankton respiration in the mesopelagic zone. They calculated active transport rates ranging from 0.42 – 0.70 mmol C m^-2^ d^-1^, which equates to only 4 – 14% of the sinking carbon export that we measured at the base of the euphotic zone. However, as most of these vertical migrants likely migrate to depths >400 m during the day, this may be equal to 18 – 35% of the sinking carbon that reached our deep sediment traps at ∼440 m. For the GoM, Kelly et al. (2021) estimated that active transport was equal to a similar ∼10% of sinking particle export flux at the base of the euphotic zone. These modest estimates of active transport are similar to those found in other oligotrophic regions. For instance, at the Hawaii Ocean Time-series site, vertical migrants were responsible for 19% of the carbon exported as sinking particles (Hannides *et al*., 2009), while at the Bermuda Atlantic Time-series site they were responsible for 8% (Steinberg et al., 2000). These results point to an emerging view that active transport is a minor, but not negligible component of export flux in low-productivity regions (Archibald *et al*., 2019; Nowicki et al., 2022; Stukel et al., 2022a), although the deep depth of respiratory C loss for vertical migrants can lead to disproportionately high carbon sequestration (Bianchi *et al*., 2013; Boyd et al., 2019; Stukel et al., 2023).

### 4.2. Particle flux and the functioning of the deep chlorophyll maximum

Sediment traps are seldom deployed in the euphotic zone. One reason for this is that hydrodynamic biases can be especially pronounced during turbulent conditions in the mixed layer (Baker *et al*., 1988; Buesseler *et al*., 2007). This issue is unlikely to be problematic, however, in strongly stratified conditions with shallow mixed layers as encountered on the BLOOFINZ-GoM and BLOOFINZ-IO cruises, and indeed in many areas with prominent DCMs. Another reason that sediment traps are usually only deployed beneath the euphotic zone is that researchers often define export flux as the flux leaving the base of the euphotic zone or a specific depth horizon (often 100 m) and are mainly interested in variability in this export flux as well as remineralization at deeper depths (Henson et al., 2011; Marsay et al., 2015; Palevsky and Doney, 2018; Buesseler *et al*., 2020). While export flux and flux attenuation are important biogeochemical processes to study, our results suggest that shallower sediment trap deployments can illuminate interesting patterns that may help in understanding the dynamics and vertical structure of the euphotic zone.

Our results show surprisingly high export flux from the upper euphotic zone to the lower euphotic zone (Fig. 8, with the upper and lower euphotic zone defined operationally based on the depth of deployment of our sediment traps, but approximately dividing the euphotic zone equally). In the Argo Basin, sinking particle flux at a depth of ∼60-m averaged 8.1 mmol C m^-2^ d^-1^, which would equate to 27% of the NPP above this depth horizon. In the more oligotrophic Gulf of Mexico, the proportion of upper euphotic zone NPP exported was even higher, with a flux of 6.4 mmol C m^-2^ d^-1^ equating to 36% of upper euphotic zone NPP. These results for the Gulf of Mexico are also supported by ^238^U-^234^Th disequilibrium results, which suggest that sinking particle flux above the DCM averaged 3.5 mmol C m^-2^ d^-1^ and sinking particle flux beneath the euphotic zone averaged 2.7 mmol C m^-2^ d^-1^ (Stukel et al., 2021). In both regions, the sinking carbon flux from the upper to the lower euphotic zone was approximately equal to the total NPP of the lower euphotic zone. In the Argo Basin 8.1 mmol C m^-2^ d^-1^ sank, compared to lower euphotic zone NPP of 7.8 mmol C m^-2^ d^-1^, while in the Gulf of Mexico 6.4 mmol C m^-2^ d^-1^ sank and lower euphotic zone NPP was 7.9 mmol C m^-2^ d^-1^. This suggests that sinking carbon flux must be a quantitatively important carbon input to the lower euphotic zone. Distinct shifts in pigment composition of sinking material (from chlorophyll to phaeopigments) further shows that the lower euphotic zone is a depth stratum in which sinking particles are consumed and re-packaged before they exit the base of the euphotic zone. Using carbon:Chl*a* estimates for the upper euphotic zone (Selph et al., 2021; Yingling et al., this issue), we can also get an estimate of phytoplankton carbon flux into the lower euphotic zone. These estimates suggest that 0.74 mmol phytoplankton C m^-2^ d^-1^ exited the upper euphotic zone in the Argo Basin and 0.25 mmol phytoplankton C m^-2^ d^-1^ exited the upper euphotic zone in the GoM. This sinking phytoplankton flux is modest (<10%) relative to the NPP of the lower euphotic zone. However, even this relatively small flux of phytoplankton may be important when compared to the net growth of phytoplankton in the deep chlorophyll maximum, where protistan grazing often balances (or even exceeds) phytoplankton growth rates, yielding net growth rates <0.1 d^-1^ (Landry *et al*., 2008; Selph *et al*., 2016; Landry et al., 2021; Landry *et al*., this issue-b). The importance of this sinking flux may also be greater when focusing only on the dynamics of large phytoplankton. Although our measurements do not allow us to determine which phytoplankton taxa contributed to the measured chlorophyll flux, it is notable that *Prochlorococcus*, which dominated the phytoplankton biomass during BLOOFINZ-IO and BLOOFINZ-GoM (Selph et al., 2021; Yingling et al., this issue), is too small to sink as individual cells. Instead, it is likely that they and other picophytoplankton were concentrated onto appendicularian houses or aggregated into other marine snow prior to sinking. This flux of large, but potentially slowly sinking aggregates (as suggested by UVP6 results), may have been an important nutritional subsidy for mesozooplankton residing in the deep euphotic zone. Indeed, a relatively large component of the mesozooplankton community were grasping, particle-associated copepods including *Oncaea* and *Corycaeus* (Swalethorp et al., this issue), suggesting that this sinking carbon may have implications for food-web transfer in the lower euphotic zone.

**Figure 8.**
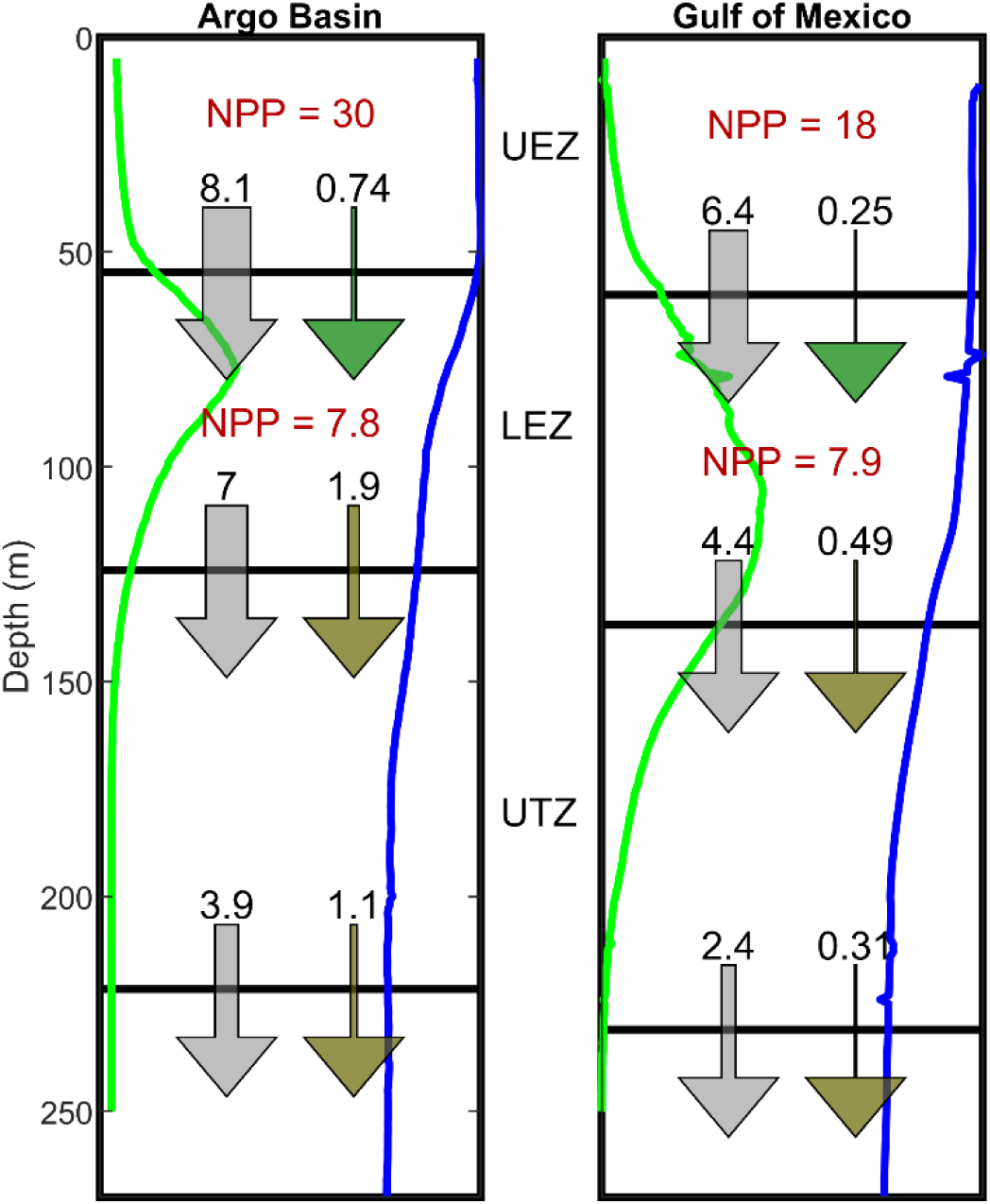
Average carbon budget for upper and lower euphotic zones (UEZ and LEZ, respectively) and upper twilight zone (UTZ) in the Argo Basin (left) and Gulf of Mexico (right). Gray arrows are sinking particle flux (mmol C m^-2^ d^-1^). Green arrows are sinking phytoplankton flux from the upper euphotic zone to the lower euphotic zone (mmol C m^-2^ d^-1^). Brown arrows are sinking herbivore fecal pellet flux (mmol C m^-2^ d^-1^). Red numbers are net primary production (mmol C m^-2^ d^-1^). Green profiles are fluorescence and blue profiles are oxygen. Profiles are only shown to highlight vertical structure. For actual values, see Fig. 1 for Argo Basin and Stukel et al. (2021) for Gulf of Mexico.

Our results also suggest that the lower euphotic zone is a region of net particle remineralization. In the Argo Basin, mean sinking carbon flux at the base of the euphotic zone (0.1% light level) was on average only 86% of sinking particle flux leaving the upper euphotic zone. In the Gulf of Mexico, the decrease was even stronger with only 69% of upper euphotic zone export flux exiting the euphotic zone. This implies that the DCM, which has substantial phytoplankton biomass, is nevertheless net heterotrophic. Another way that we can address the question of net heterotrophy vs. net autotrophy is by investigating oxygen profiles. If we make the simplifying assumption that horizontal advection and diffusion are negligible to the water column oxygen budget, then the rate of O_2_ change follows the equation:

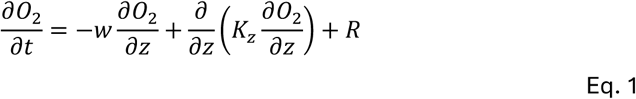

where *w* is the vertical velocity, *K_z_* is the vertical eddy diffusivity, and *R* is the net biological production or consumption of oxygen (O_2_). Vertical velocities are typically weakly downward in oligotrophic regions and this fact, combined with decreasing oxygen with depth near the DCM (Supp. Fig. 4) suggests that vertical advection is a net supply of oxygen to the DCM. Thus, if the advection term is neglected, heterotrophy is underestimated. If we further assume a quasi-steady state and that vertical eddy diffusivity is constant with depth, Eq. 1 becomes:

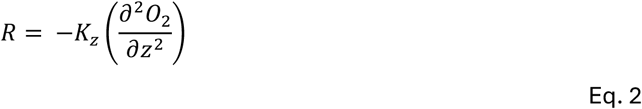

This equation suggests that the community will be net heterotrophic when the second derivative of oxygen is positive. If we integrate this second derivative over the depths of the lower euphotic zone (as defined in Fig. 8), we find that it is slightly positive (Supp. Fig. 4d) and thus agrees with the hypothesis that the deep euphotic zone is net heterotrophic. Investigating the vertical structure also shows that the second derivative switches from negative to positive at almost exactly the depth of the DCM, which would suggest that the water column switches from net autotrophic to net heterotrophic right at the DCM. We caution, however, that our assumption of no vertical advection likely leads to an underestimation of heterotrophy in the system (as a result of the likely downward average velocities noted above). This would suggest that the switch to heterotrophy might occur at slightly shallower depths than predicted by the second derivative of oxygen. Nevertheless, it is clear that particle consumption and repackaging and net remineralization can begin within the deep euphotic zone and likely shapes the carbon balance of the DCM as well as the characteristics of sinking particles exiting the euphotic zone.

### 3.5. Conclusions

The Argo Basin is a classic tropical 2-layer stratified, oligotrophic-ocean ecosystem with a low-nutrient low-chlorophyll upper layer and with a strong DCM and nitracline in the lower layer. Sinking organic carbon flux is commensurately low at the base of the euphotic zone, but export efficiency is moderate with EZ-ratios ranging from 0.085 to 0.23, similar to those in the slightly more oligotrophic central Gulf of Mexico. In both of these regions, mean carbon flux out of the upper euphotic zone (i.e., 50 – 60 m) was higher than particulate carbon export from the base of the euphotic zone (0.1% light level), suggesting that the deep euphotic zone and DCM is a zone of net remineralization. A distinct shift in the characteristics of sinking particles was also noted between these depth horizons, with sinking chlorophyll flux suggesting that intact phytoplankton were an important component of sinking carbon flux from the upper to lower euphotic zone, while phaeopigments indicated that fecal pellets of herbivorous zooplankton were more important beneath the euphotic zone. Within the twilight zone, flux attenuation was fairly typical for a marine ecosystem, with an average of 62% of sinking carbon flux penetrating an additional 100 m beneath the euphotic zone in the Argo Basin and 52% in the Gulf of Mexico. Taken together, our results suggest that export efficiency is greater in these two marginal seas than is typical in open-ocean systems that are equally oligotrophic in terms of other indices of trophic state.

## CRediT authorship contribution statement

**Michael R. Stukel:** Conceptualization, Funding acquisition, Investigation, Writing – original draft, **Tristan Biard:** Data curation, Resources, Writing – review and editing, **Moira Decima:** Conceptualization, Investigation, Writing – review and editing, **Christian K. Fender:** Investigation, Writing – review and editing, **Opeyemi Kehinde:** Investigation, Writing – review and editing, **Thomas B. Kelly:** Investigation, Writing – review and editing, **Sven A. Kranz:** Conceptualization, Funding acquisition, Investigation, Writing – review and editing, **Manon Laget:** Investigation, Writing – review and editing, **Michael R. Landry:** Conceptualization, Funding acquisition, Investigation, Writing – review and editing, **Natalia Yingling:** Investigation, Writing –review and editing

## Acknowledgments

We thank our many collaborators in the BLOOFINZ-IO and BLOOFINZ-GoM research projects and are very grateful to the captain and crew of the R/V Roger Revelle. This study was supported by National Science Foundation biological oceanography grants OCE-1851347 to M.R.S. and S.A.K. and OCE-1851558 to M.R.L. This study was a part of project EP-46 of the 2nd International Indian Ocean Expedition (IIOE-2). Sampling in the Argo-Rowley Terrace Marine Park was done under Australian Government permit AU-COM2021-520 and permit PA2021-00062-1 issued by the Director of National Parks, Australia.

## Data availability

Data from this project are available at the Biological and Chemical Oceanography Data Management Office (BCO-DMO) project pages for BLOOFINZ-IO and BLOOFINZ-GoM: https://www.bco-dmo.org/project/819488 and https://www.bco-dmo.org/project/834957

## Supplementary Figures

**Supp. Fig. 1.**
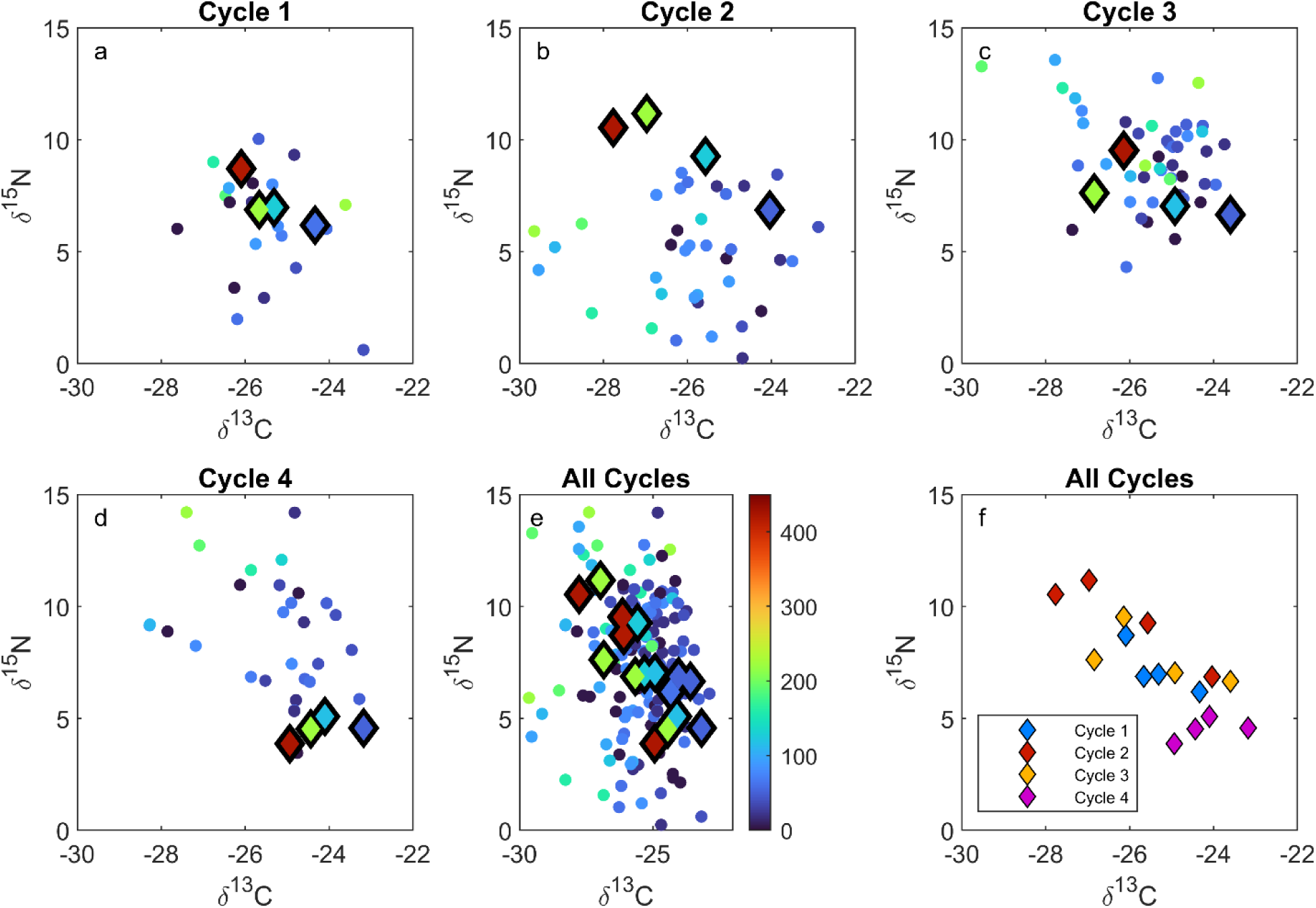
Carbon and nitrogen isotopic ratios for seston (small circles) and sinking particles (larger diamonds) for a) Cycle 1, b) Cycle 2, c) Cycle 3, d) Cycle 4, e) All cycles combined, and f) Sediment trap data only from all cycles combined. Color indicates depth in all panels except panel f and corresponds to the colorbar shown in panel e.

**Supp. Fig. 2.**
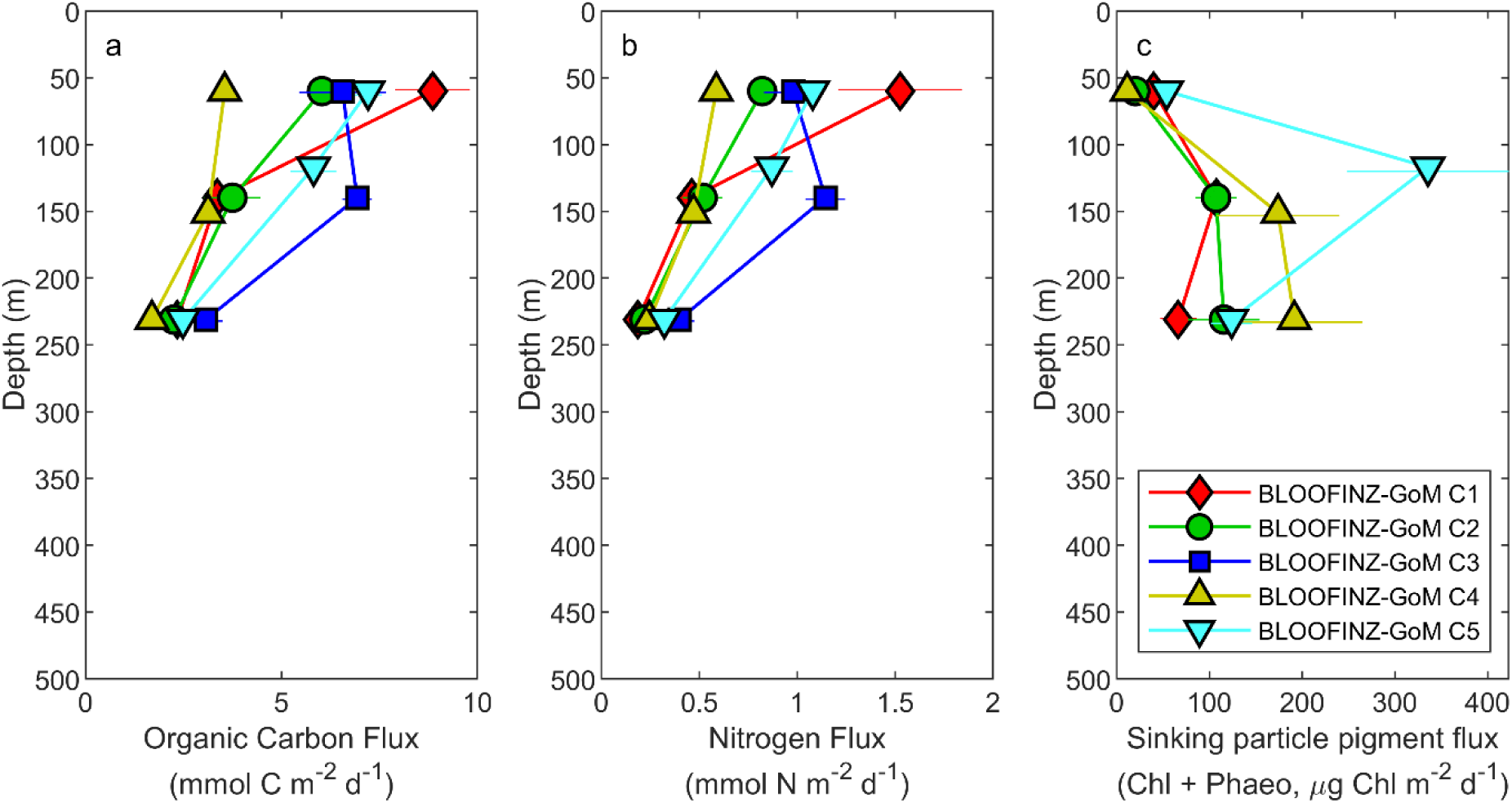
Sinking particle organic carbon flux (a), nitrogen flux (b) and pigment flux (c) on the BLOOFINZ-GoM cruise (compare to BLOOFINZ-IO data presented in Fig. 5). Data is from Stukel et al. (2021).

**Supp. Fig. 3.**
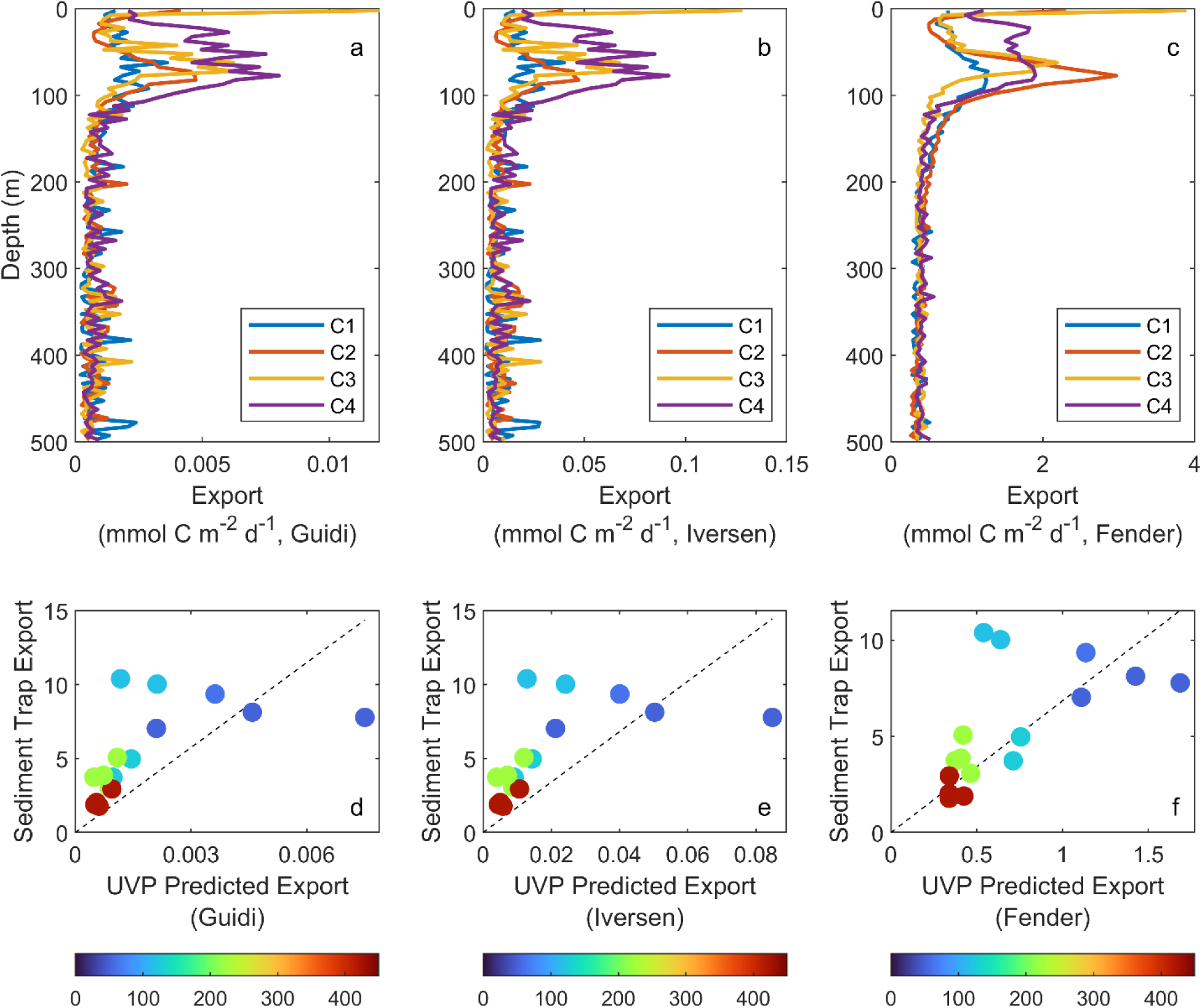
Estimated sinking particle carbon export from particle-size spectra from the Underwater Video Profiler, calculated using the equation: 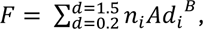 where *n_i_* is the concentration of particles of size class *i* and *d_i_* is the equivalent spherical diameter (mm) of particles of size class *i*, and *F* is sinking carbon flux (mg C m^-2^ d^-1^). A and B are constants calculated in prior studies. (a) and (d) use A = 12.5 and B = 3.81 from Guidi et al. (2008). (b) and (e) use A = 273.8 and B = 4.27 from Iversen et al. (2010). (c) and (f) use A = 15.4 and B = 1.05 from Fender et al. (2019). (a) – (c) show vertical profiles of UVP-estimated export flux. (d) – (f) show comparisons of UVP-estimated export flux (x-axis) at the depth of sediment trap carbon export measurements (y-axis). Dashed line is a linear regression forced through the origin. Points above the dashed line show where actual sinking particle flux was relatively high compared to the estimate from the particle size spectrum (i.e., particles were sinking faster than predicted by their size). Points below the dashed line represent actual sinking particle flux that was relatively low compared to the estimate of the particle size spectrum. We note that previous results suggest that the particle size spectrum should not be expected to provide sufficient information for estimating the absolute magnitude of flux. Rather, we present this data as a way of visualizing relative patterns in export flux.

**Supp. Fig. 4.**
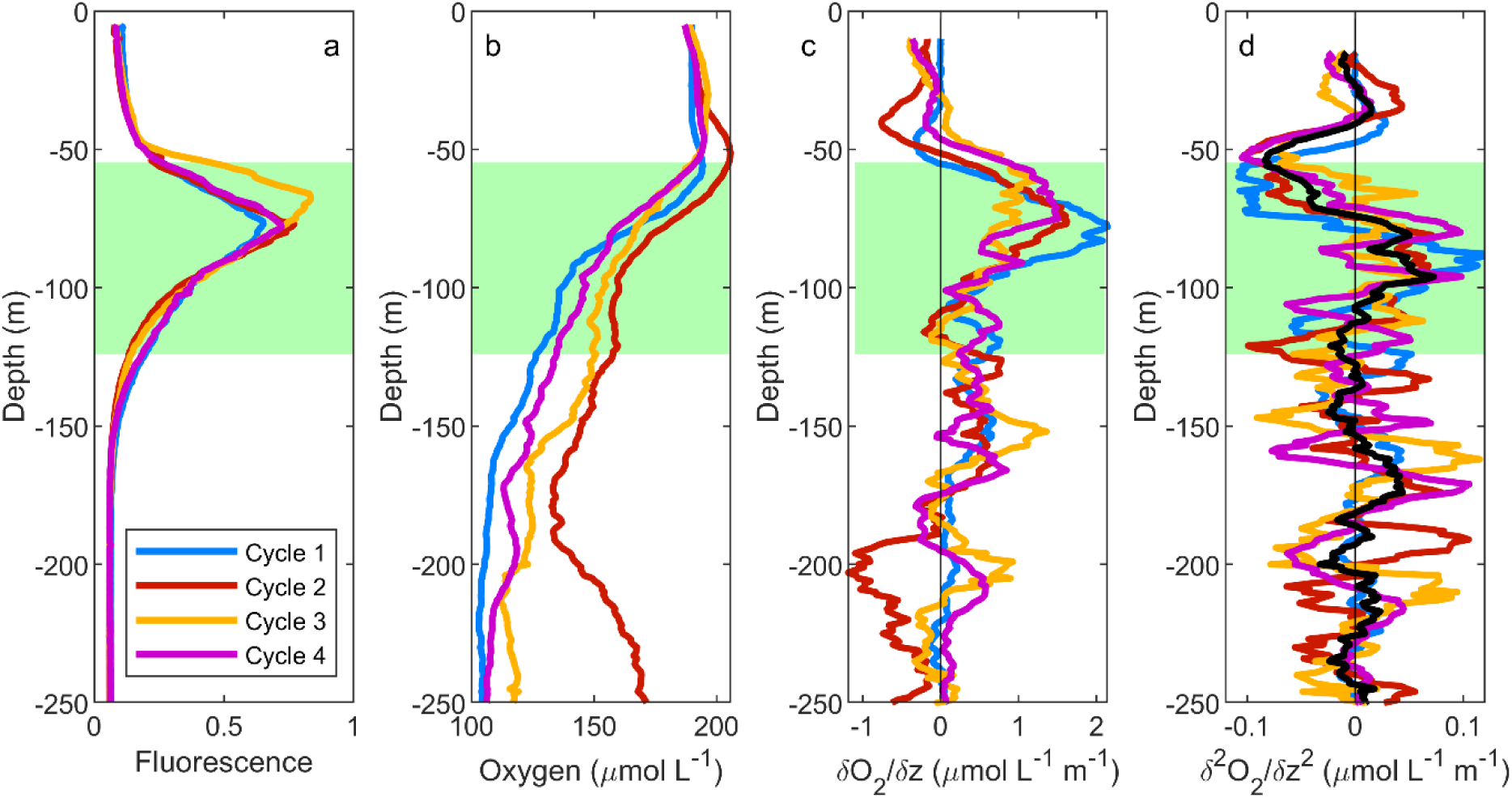
Oxygen dynamics during BLOOFINZ-IO. (a) Cycle average vertical profiles of fluorescence for comparison. Green boxes in each panel show the extent of the lower euphotic zone as defined as the depth range between our shallowest sediment trap and the base of the euphotic zone (i.e., same depth ranges as shown in Fig. 8). (b) Cycle average vertical oxygen profiles. (c) Derivative (with respect to depth) of oxygen. Oxygen profile was smoothed with a 10-m running mean before calculating derivate. (d) Second derivate (with respect to depth) of oxygen. Derivative in (c) was smoothed with a 10-m running mean before calculating second derivative in (d).

